# AptamerRunner: An accessible aptamer structure prediction and clustering algorithm for visualization of selected aptamers

**DOI:** 10.1101/2023.11.13.566453

**Authors:** Dario Ruiz-Ciancio, Suresh Veeramani, Eric Embree, Chris Ortman, Kristina W. Thiel, William H Thiel

## Abstract

Aptamers are short single-stranded DNA or RNA molecules with high affinity and specificity for targets and are generated using the iterative Systematic Evolution of Ligands by EXponential enrichment (SELEX) process. Next-generation sequencing (NGS) revolutionized aptamer selections by allowing a more comprehensive analysis of SELEX-enriched aptamers as compared to Sanger sequencing. The current challenge with aptamer NGS datasets is identifying a diverse cohort of candidate aptamers with the highest likelihood of successful experimental validation. Herein we present AptamerRunner, an aptamer clustering algorithm that generates visual networks of aptamers that are related by sequence and/or structure. These networks can then be overlayed with ranking data, such as fold enrichment or data from scoring algorithms. The ability to visually integrate data using AptamerRunner represents a significant advancement over existing clustering tools by providing a natural context to depict groups of aptamers from which ranked or scored candidates can be chosen for experimental validation. The inherent flexibility, user-friendly design, and prospects for future enhancements with AptamerRunner has broad-reaching implications for aptamer researchers across a wide range of disciplines.

**GRAPHICAL ABSTRACT:** 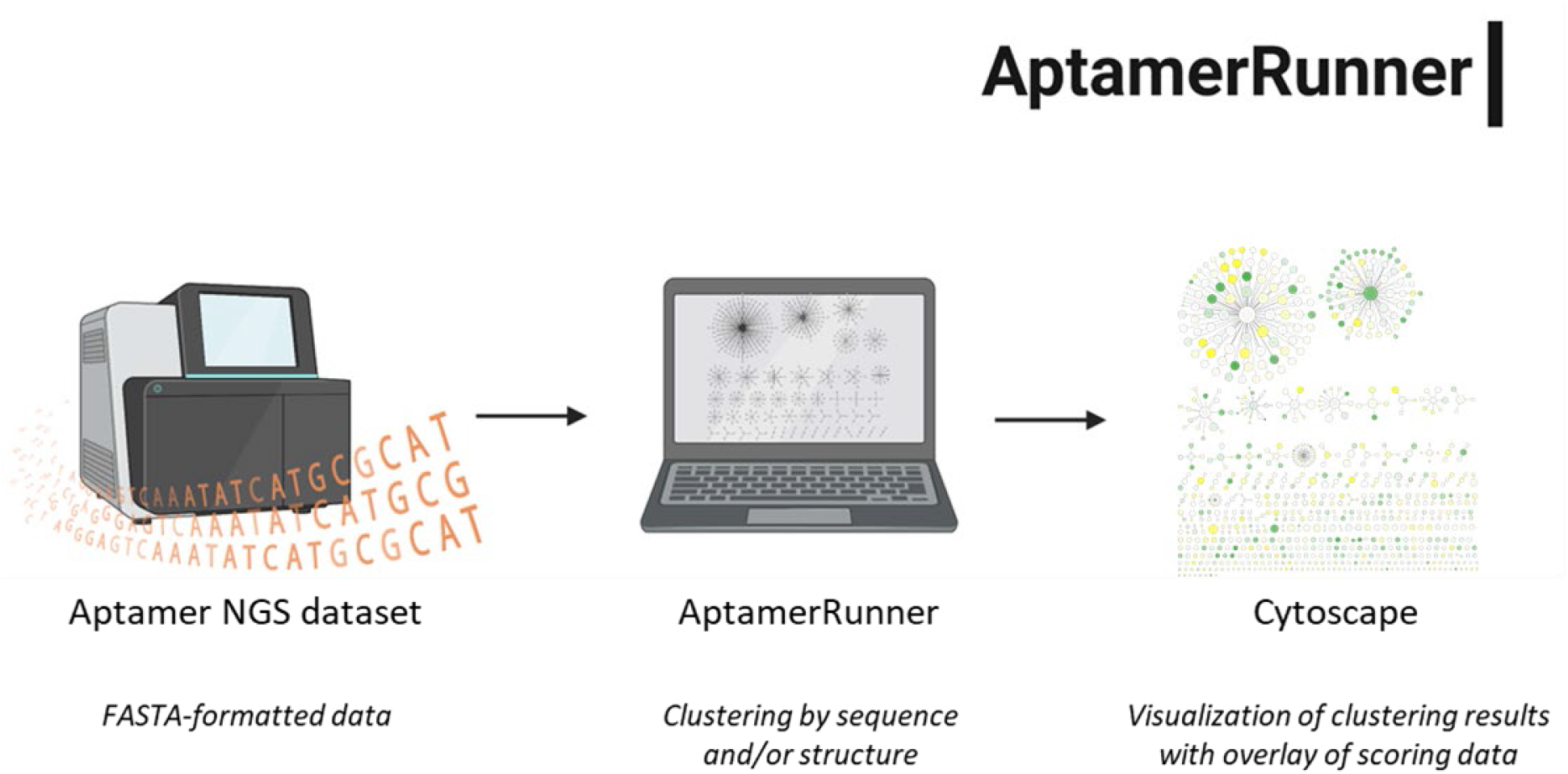

## INTRODUCTION

Aptamers are short synthetic RNA or DNA oligonucleotides that recognize target epitopes with specificity and affinity analogous to antibody-antigen interactions (1). Applications of aptamers are broad-reaching and include biosensors (2), research reagents (3,4), tools for mechanistic discovery (3), diagnostics (5), and therapeutics (5). Recently the aptamer avacincaptad pegol, that target complement C5, was FDA-approved for the treatment of geographic atrophy (6). Aptamers are generated using an *in vitro* process known as Systematic Evolution of Ligands by EXponential enrichment (SELEX) (7,8). The SELEX process now includes numerous variations with new SELEX strategies constantly being developed (1). At the completion of SELEX, selected aptamers are identified by sequencing, with the field shifting from Sanger sequencing towards next-generation sequencing (NGS) platforms. NGS yields hundreds of millions of reads, with each read containing the entirety of an aptamer sequence. Thus, the aptamers enriched during SELEX can now be interrogated to a degree not achievable with Sanger sequencing (9,10). However, these large NGS datasets have created new challenges in terms of how parse the millions of reads to identify the best candidate aptamers to then validate experimentally. Testing thousands or even hundreds of aptamer candidates is still unattainable for most aptamer researchers; therefore, identification of the top candidates is paramount. To address this need, several bioinformatics approaches specific for aptamer NGS datasets have emerged (11,12). The analysis of an aptamer NGS dataset includes processing the FASTQ data and application of strategies to identify candidate aptamers using various motif identification, scoring, and clustering algorithms (11,12). Bioinformatics tools to identify candidate aptamers are frequently applied in concert, for example clustering is used to identify separate groups of aptamers (13–15), followed by ranking aptamers within these groups using fold enrichment or scoring algorithms (16,17).

The central theory behind aptamer clustering is that aptamers that are closely related based on their sequence and predicted structures are likely to target the same epitope (11,12). The earliest efforts to cluster aptamers used sequence alignment algorithms such as ClustalW (18–20), but these data can be difficult to interpret and this method does not take into account the predicted structures of the aptamers. Our group introduced the concept of clustering aptamers by either sequence relatedness using Levenshtein edit distance or by predicted secondary structure relatedness using tree distance (15). We applied a clustering strategy, termed the *all-vs-all* approach, wherein all aptamers within a dataset are compared to each other. Clusters were defined as the aptamers that interconnect within a threshold distance measure (e.g., edit distance of 3). However, this early clustering algorithm was not easily accessible and thus has not been widely adopted. The current prevailing clustering algorithms include AptaCluster (21,22), FASTAptamer (23), and FASTAptameR 2.0 (24). These tools generate networks of related aptamers using either Hamming or Levenshtein edit distance and introduced a new concept for clustering aptamers termed the *seed approach*. The seed approach generates networks of aptamers centered around a *seed sequence*, which is defined as the most abundant sequence within an aptamer NGS dataset. All aptamers that connect to the seed sequence within a threshold edit distance measure are designated as a cluster. This process is iterated by removing the initial seed sequence and all connected aptamer sequences from further analysis, and the next most abundant sequence becomes the seed for the next cluster. These seed-based aptamer clustering algorithms are significantly more accessible than our initial algorithm, and the seed approach has significant computational advantages over the all-vs-all approach. A limitation of the seed approach is that it is a greedy process that can potentially miss important inter-aptamer relationships identified by the all-vs-all clustering approach. In addition, the available seed-based algorithms do not consider structure when generating networks of related aptamers. The output from these algorithms is text-based rather than graphically represented, which severely limits interpretation of the clustering results and prevents integration of ranking data that is necessary to identify candidates within the groups of interconnected aptamers.

Herein, we introduce *AptamerRunner*, an accessible aptamer structure prediction and clustering algorithm for the visualization of networked aptamers. AptamerRunner was designed with a focus on flexibility to ensure that aptamer researchers can apply different clustering strategies to suit their needs. AptamerRunner includes several novel clustering features not previously available: 1) the option to use either the all-vs-all approach or the seed approach; 2) the option to interrogate both sequence and structure relatedness simultaneously by applying logical operators (AND, OR) with edit and tree distance thresholds; and 3) inclusion of distance measure data as metadata within the edges of the interconnected aptamer sequences. To provide easier access and functionality, AptamerRunner and all dependencies have been packaged into a Docker image (25) that is operated by command line using a platform-independent .NET script. The AptamerRunner clustering algorithm outputs results as an eXtensible Graph Markup and Modeling Language (XGMML) file that permits graphical representation of the clustering results using network analysis programs such as the open-source Cytoscape program (26). Importantly, through the graphical representation of the clustering data, additional information such as ranking data can be overlayed onto the networked aptamers to facilitate an integrated analysis whereby clusters of aptamers and ranking data can be interpreted at the same time. This integration of clustering with ranking data presents a novel analysis of selected aptamers that aids in the identification of the best candidates.

## MATERIALS AND METHODS

### AptamerRunner

AptamerRunner is a .NET program coded in C# that generates the Docker commands to adapt to various operating system constraints. AptamerRunner will check the Docker repository and download the most recent AptamerRunner Docker image (for detailed instructions to use AptamerRunner refer to supplemental methods). The AptamerRunner Docker image is comprised of two independent python algorithm components (**Figure 1**). The first component predicts the secondary structure of RNA or DNA aptamer sequences (**Figure 1A**). The second component calculates aptamer relatedness to generate networks of related aptamer sequences (**Figure 1B**). This segmentation permits each component to be implemented independently and provides additional flexibility to the user. Once the AptamerRunner Docker container has completed running the structure predication algorithm or clustering algorithm, AptamerRunner will close the AptamerRunner Docker container. The AptamerRunner .NET script is available at https://github.com/ui-icts/aptamer-runner/releases/tag/v0.0.3 and within supplemental data.

**Figure 1.**
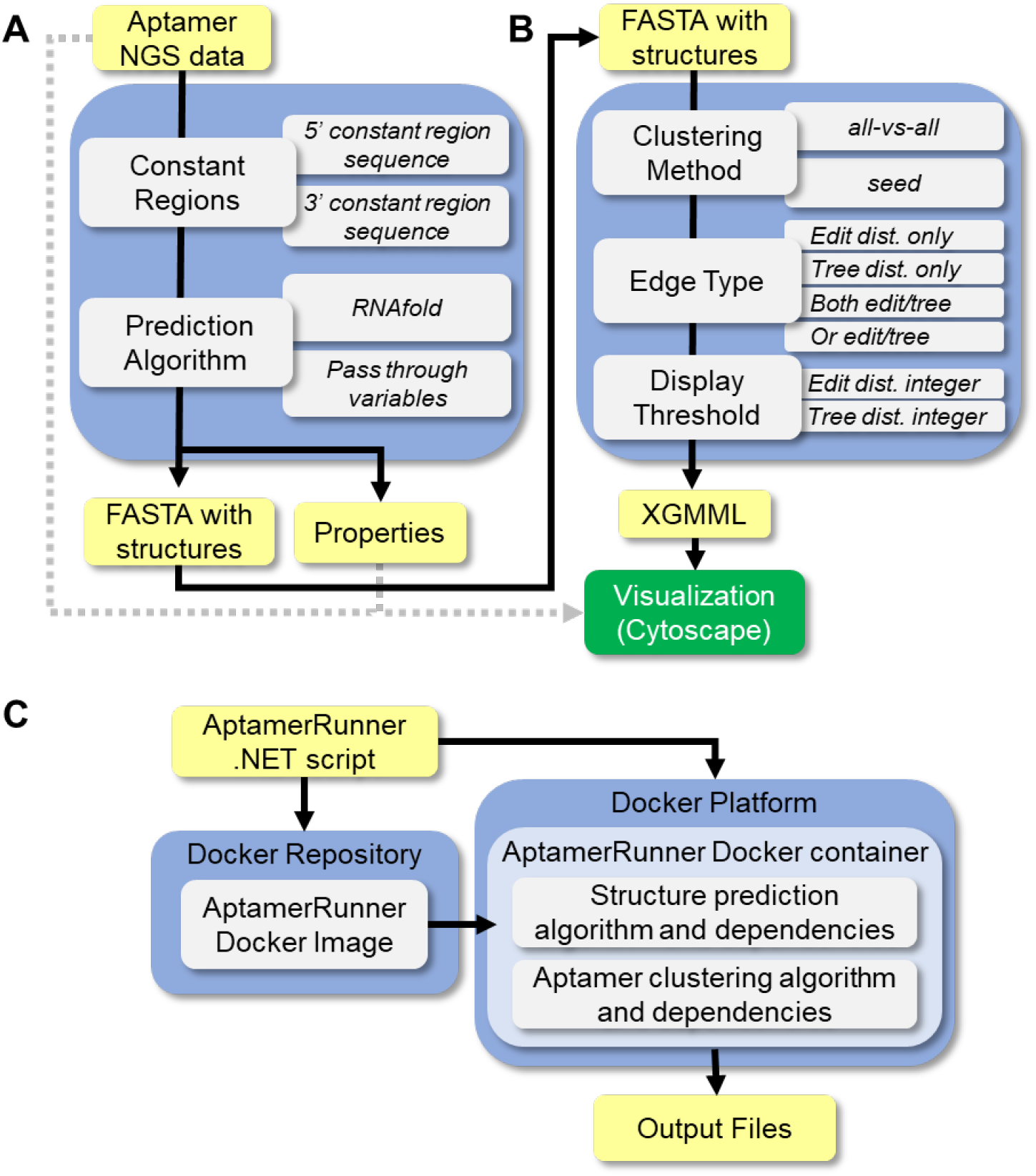
AptamerRunner, a structure prediction and clustering algorithm a to visualize selected aptamers. AptamerRunner consist of two independent algorithms, a secondary structure prediction algorithm and an aptamer clustering algorithm. (**A**) The secondary structure prediction algorithm utilizes collapsed aptamer NGS data in FASTA format to predict the secondary structure of a full-length aptamer using RNAfold. The secondary structure prediction algorithm has the option to append the constant region sequence if needed. Output includes a modified FASTA-formatted file with a third line containing the dot-bracket annotation of each predicted structure and a properties file that includes information about the predicted structures (e.g., minimum free energy). (**B**) The AptamerRunner clustering algorithm uses the FASTA-formatted file with the predicted structures to generate networks of related aptamers using options selected by the user for the *Clustering Method*, the *Edge Type,* and the *Display Threshold*. Output files include the clustering results compiled into a XGMML file, which is visualized using Cytoscape, a log file, and the input file. (**C**) AptamerRunner is a .NET bash script coded in C#. The AptamerRunner .NET script communicates with the Docker repository to download the latest version of the AptamerRunner Docker image and then initiates the AptamerRunner Docker image within the Docker platform as a Docker container. The Docker container includes the AptamerRunner structure prediction algorithm and the clustering algorithm with all dependencies. Once either algorithm has finished processing any user commands and output results, the AptamerRunner .NET script shuts down the AptamerRunner Docker container.

### RNA or DNA aptamer secondary structure prediction algorithm

The structure prediction component (**Figure 1A**) of AptamerRunner requires either full-length aptamers or only the variable region of the aptamers in a FASTA-formatted file. The FASTA file should contain collapsed aptamer NGS data, in which all unique aptamer sequences are represented once and are ranked in descending order based on number of duplicate reads (**Figure S1A**). If the FASTA file contains only the variable region sequences, AptamerRunner provides an option to automatically append the constant regions per user input. Secondary structures are predicted using the RNAfold structure prediction algorithm from the Vienna Package v2.0 (27–29), with the lowest minimum free energy structure being retained. Pass-through commands specific for RNAfold can be included during AptamerRunner execution. The predicted aptamer secondary structures are appended to a modified FASTA-formatted file (FASTA.struct) using dot-bracket annotation (**Figure S1**). Properties associated with the predicted structures (e.g,. minimum free energy) that are output by the RNAfold are compiled into a separate tab-delimited file with the aptamer FASTA header information as a key. These structural properties of each aptamer sequence can be imported, if needed, for overlaying when visualizing the clustering data.

### Aptamer clustering algorithm

The aptamer clustering component of AptamerRunner generates networks of related aptamers from aptamer sequences and predicted secondary structures (**Figure 1B**). Networks of related aptamers are constructed using either the all-vs-all approach or the seed approach. User options include selecting the 1) *Edge Type* and the 2) *Maximum Distance Measure*.

#### Edge Type

The Edge Type defines the relatedness distance metric applied by the clustering algorithm in order to build networks of connected aptamers. Sequence similarity is determined using Levenshtein edit distance (30) and structure similarity is determined using tree distance (31). These distance measurements are calculated by the clustering algorithm using RNAdistance from the Vienna Package v2.0 (27,32). Options for clustering include 1) *edit* to use only edit distance data, 2) *tree* for tree distance data only, 3) *both* to apply the logical operator AND with edit distance and tree distance data, and 4) *or* to apply the logical operator OR with edit and tree distance data. Regardless of Edge Type applied, AptamerRunner will calculate both the edit and tree distances between aptamer sequences and include these data as metadata for the edges connecting two aptamer sequences (**Figure S2**). This enables users to perform additional analyses using the edit and tree distance values within Cytoscape. For example, if *edit* is designated as the Edge Type option, AptamerRunner will only apply edit distance data when constructing the networks of related aptamers, but it will also calculate tree distance values and include these data as edge metadata.

#### Maximum Distance Measure

The Maximum Distance Measure determines the maximum edit and/or tree distance value for a given Edge Type from which networks of related aptamers will be constructed. A separate Maximum Distance Measure can be set for edit distance and for tree distance. The clustering algorithm requires that the Maximum Distance Measure match the Edge Type and, if using a logical operator, that a Maximum Distance Measure be set for both edit distance and tree distance.

#### Output Files

Three files are outputted by the clustering program: 1) an eXtensible Graph Markup and Modeling Language (XGMML) file containing the aptamer clustering data, which can be imported into Cytoscape; 2) the input FASTA.struct file; and 3) a log file of the commands used to initiate AptamerRunner.

### Cytoscape visualization of AptamerRunner clustering results

The XGMML file of clustering results from AptamerRunner can be visualized through Cytoscape (26), an open-source network analysis program. Networks of clustered aptamer sequences are visualized using *nodes*, which represent unique aptamer sequences, and *edges* that connect related aptamer sequence nodes (**Figure S2**). Nodes and edges include metadata associated to each aptamer sequence (e.g., name, sequence, structure) and between interconnected aptamers (e.g., edit distance and tree distance values). Additional metadata, such as normalized read counts, fold enrichment or data from scoring algorithms, can be imported into Cytoscape’s node data tables from comma or tab delimited text files. These metadata can be used to dictate the visual properties of nodes and edges within Cytoscape to facilitate interpretation of the clustering results. Cystoscope’s interactive interface permits easy selection of nodes to isolate potential candidate aptamer sequences. Nodes selected from networks are compiled into Cytoscape’s data table along with corresponding node data (**Figure S3A**). The Cytoscape data table may be copied directly or exported as a text file. In addition to selecting individual aptamer sequences, groups of aptamers may be readily identified and examined in more detail using network analysis tools such as the clusterMaker algorithm (33). The clusterMaker algorithm assigns an identification to each group of related aptamers and appends these identifications to the Cytoscape data table (**Figure S3B**).

## RESULTS

To demonstrate the utility of AptamerRunner and compare to other aptamer clustering tools, we used a published aptamer NGS dataset from a selection against B-cell leukemia cells (14). With these data, we applied the various AptamerRunner clustering options with either edit distance of 1 or tree distance of 0, when applicable. When comparing AptamerRunner against FASTAptamer, FASTAptameR 2.0 and AptaCluster, we applied comparable clustering parameters using an edit distance of 1 with the seed approach.

### Visualization of AptamerRunner clustering results

A central tenet of AptamerRunner is visual representation of clustering results using nodes and edges, which provides a natural context to interpret clustering data that is significantly easier to comprehend as compared to text-based results (**Figure 2**). When the seed option is used to generate the networks of related aptamers, the seed sequence is clearly identifiable as the central node and the number of sequences connecting to each seed sequences are easily discerned (**Figure 2A** and **2B**). As compared to the seed approach, the all-vs-all option produces fewer but more complex clusters of inter-related aptamer sequences (**Figure 2C** and **2D**). The all-vs-all approach clustering results can include large, complex hairball networks (**Figure 2C**, largest cluster) and smaller clusters with naturally occurring central nodes. Importantly, clustering conducted using either the seed approach or the all-vs-all approach can provide different perspectives of the same data.

**Figure 2:**
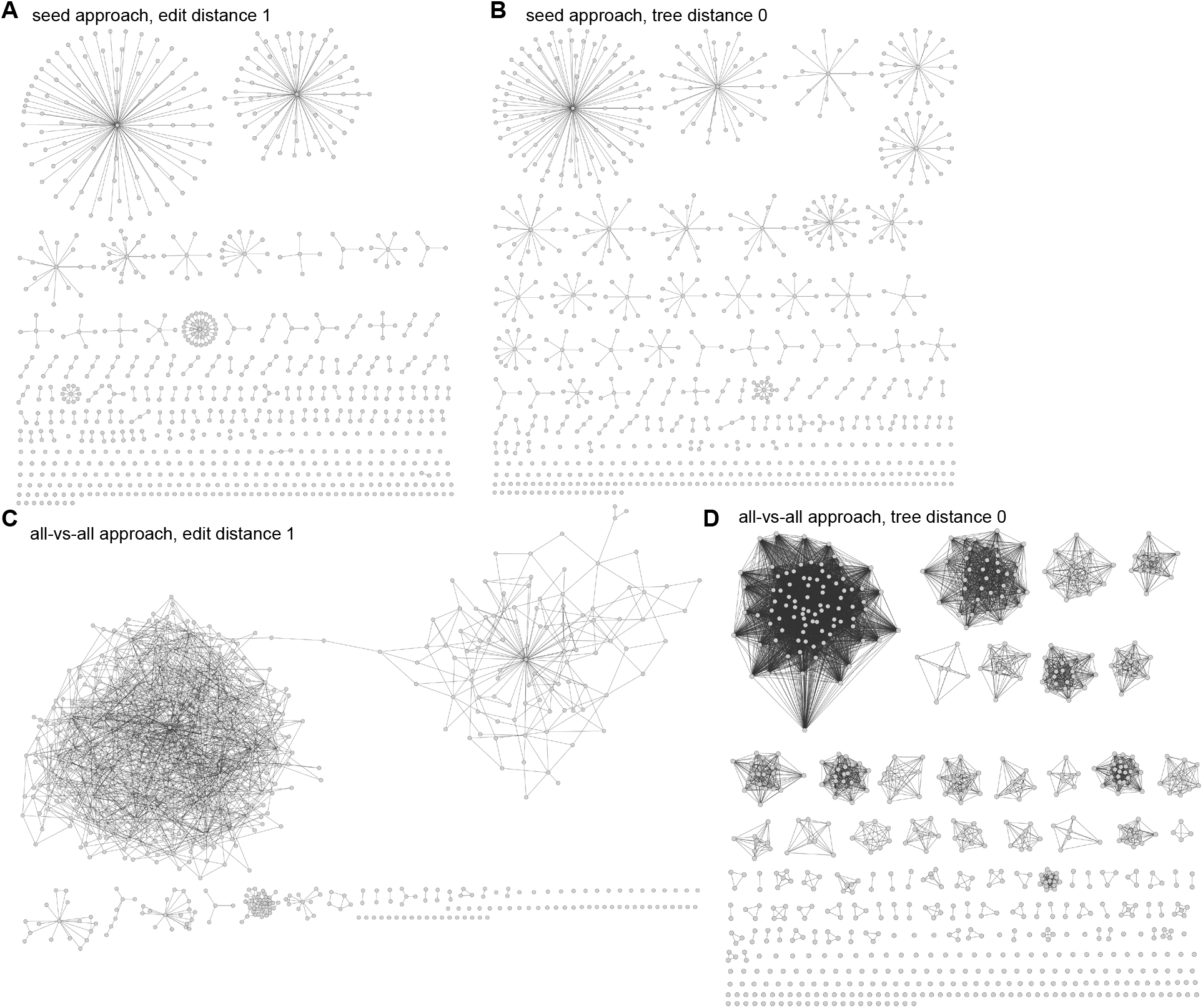
AptamerRunner all-vs-all and seed clustering approaches using either edit distance or tree distance. AptamerRunner clustering aptamer NGS data using different clustering methods with either edit distance 1 or tree distance 0. (**A**) Seed approach with edit distance 1 or (**B**) tree distance 0. (**C**) All-vs-all approach with edit distance 1 or (**D**) tree distance 0. Ruiz-Ciancio *et. al.* (14) data was used as example data for clustering and clustering results were visualized using Cytoscape (v 2.8.1).

A second important improvement within AptamerRunner as compared to other clustering algorithms is the introduction of logical operators AND and OR with edit distance and tree distance. The AND function and the OR function can be applied to gain insight into potential functional aspects of an aptamer. The AND function can reveal areas of nucleotide substitutions that are well-tolerated or that impart beneficial properties. Conversely, the OR function can reveal nucleotide changes that have a significant impact on the predicted structure of an aptamer. The use of the AND logical operator places a higher stringency, and the use of the OR logical operator less stringency, onto the networks of related aptamer generated. For example, clustering the data presented in **Figure 3** using the AND logical operator with an edit distance 1 and tree distance 0 yields smaller, more constrained networks of related aptamers when using the seed (**Figure 3A**) and all-vs-all approaches (**Figure 3B**). Conversely, the OR logical operator yields larger less constrained networks of related aptamers (**Figure 3C** and **3D**).

**Figure 3:**
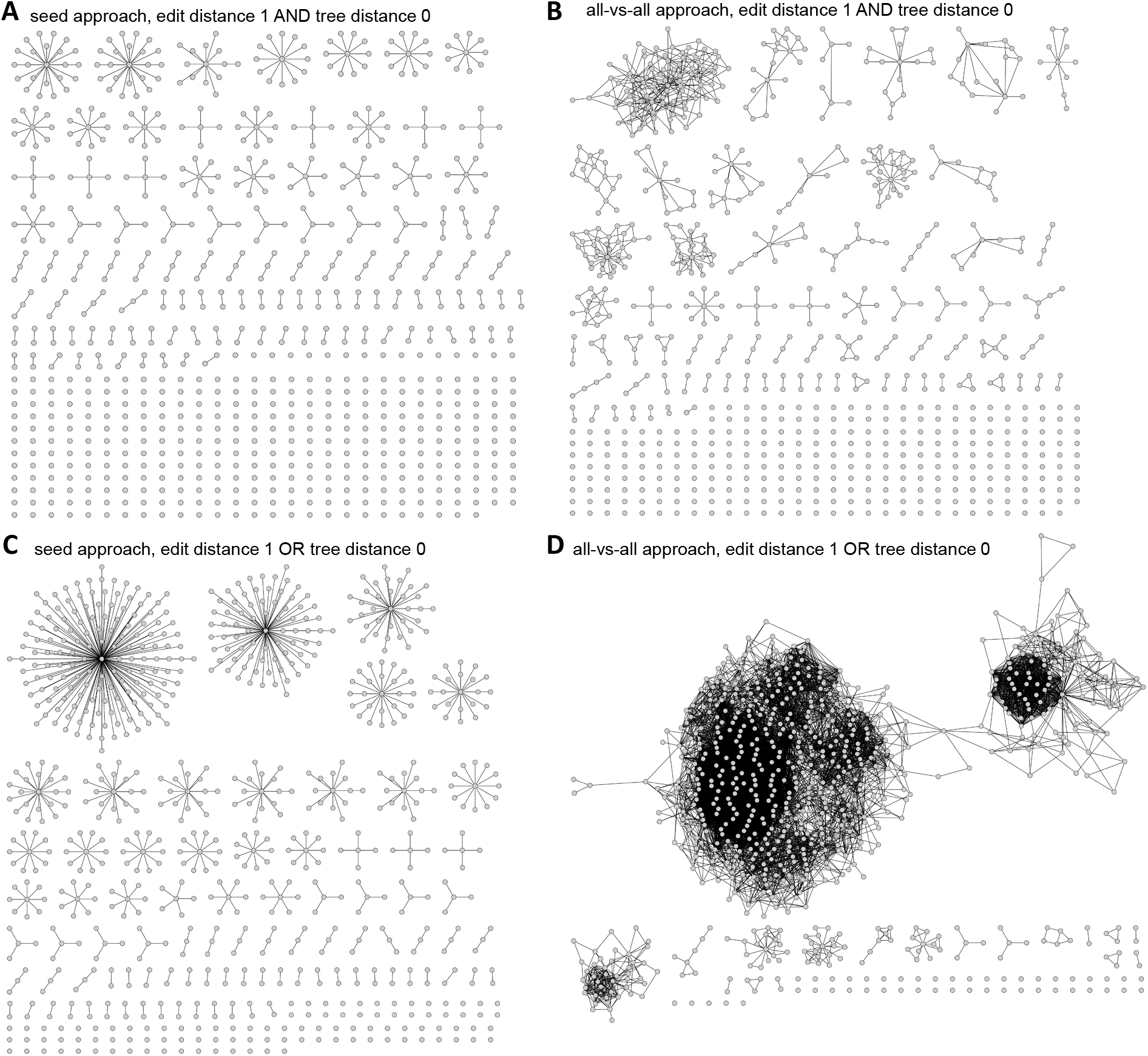
Use of logical operators (AND, OR) with AptamerRunner clustering. Clustering using logical operator AND with edit distance 1 and tree distance 0 with the (**A**) seed approach and the (**B**) all-vs-all approach. Clustering using the logical operator OR with edit distance 1 and tree distance 0 with the (**C**) seed approach and the (**D**) all-vs-all approach. Ruiz-Ciancio *et. al.* (14) data was used as example data for aptamer classification and selection, and data are visualized using Cytoscape (v 2.8.1).

### Comparison of AptamerRunner to other aptamer clustering algorithms: FASTAptamer, FASTAptameR 2.0 and AptaCluster

We next compared the capabilities and output of AptamerRunner to the capabilities and output of FASTAptamer, FASTAptameR 2.0 and AptaCluster (22-24,34). Details and features of FASTAptamer, FASTAptameR 2.0, AptaCluster and AptamerRunner are summarized in **Table 1**. The FASTAptamer clustering algorithm (*FASTAptamer-Cluster*) and FASTAptameR 2.0 clustering algorithm (*cluster module*) determine sequence families of aptamers by Levenshtein edit distance using the seed approach. FASTAptameR 2.0, an update to FASTAptamer, operates through an offline web-browser that accesses a Docker container. FASTAptameR 2.0, like FASTAptamer, applies a seed approach using edit distance only, but has a computationally faster clustering algorithm and a cluster visualization function (*cluster diversity module*) that generates a principal component plot (PCA) using a k-mer matrix of the clustered aptamer sequences. AptaCluster is provided as a component of the AptaSuite package with a GUI that is accessible as a Java application (35). AptaCluster filters input data using a local sensitivity hash function and clusters aptamers by Hamming edit distance using the seed approach.

**Table 1.**

|  | **AptamerRunner** | **FASTAptamer** | **FASTAptamer 2.0** | **AptaCluster** |
| --- | --- | --- | --- | --- |
| **Interface** | Command line C# .NET script operating a Docker container | Command line Perl-based programs | Offline web browser accessing a Docker container | AptaSuite Java application |
| **Components**<br>(*clustering) | • Secondary structure prediction<br>• Clustering algorithm* | • Count<br>• Compare<br>• Cluster*<br>• Enrich<br>• Search | • FASTAptamer components<br>• Cluster module*<br>• Edit distance module<br>• Motif discovery module<br>• Position enrich module | • AptaGUI<br>• AptaPlex<br>• AptaSim<br>• AptaCluster*<br>• AptaTrace<br>• AptaMut |
| **Third party software** | Docker, Cytoscape | none | Docker, web browser | Java |
| **Clustering approach** | seed, all-vs-all | seed | seed | seed |
| **Distance measures** | • Edit distance<br>• Tree distance<br>• Logical operators (AND, OR) | • Edit distance | • Edit distance | • Edit distance |
| **Visualization** | Cytoscape display of networked aptamers (nodes) connected by distance measures (edges) | N/A | • Cluster metaplots<br>• k-mer PCA | N/A |
| **Metadata** | Nodes:<br>• Aptamer name<br>• Predicted structure<br>• User-imported<br>Edges:<br>• Edit distance values<br>• Tree distance values<br>• User-imported | N/A | N/A | N/A |

Of note, FASTAptamer, FASTAptameR 2.0 and AptaCluster cannot perform clustering based upon predicted structures, nor do they compare all aptamers to each other regardless of their enrichment during SELEX (e.g., all-vs-all approach). AptaSuite, of which AptaCluster is a component, includes an algorithm AptaTrace that identifies sequence-structure motifs as sequence logos with secondary structure probability profiles (36). The AptaTrace algorithm does not perform clustering of these predicted secondary structure and thus is not included in the comparison to AptamerRunner. To compare AptamerRunner to FASTAptamer, FASTAptameR 2.0 and AptaCluster, we applied the seed approach using only edit distance with no logical operators, and we used aptamer NGS data from a selection against B-cell leukemia cells (14).

The FASTAptamer and FASTAptameR 2.0 clustering algorithms produced identical modified FASTA formatted text files (**Figure 4A**). Each sequence identifier within the FASTA file (denoted with the “>” annotation) contains the following data features for each aptamer sequence: sequence ID, raw read count, normalized read count, cluster ID, rank within the cluster, and edit distance from the seed sequence. For example, in **Figure 4A**, the first listed sequence has an identifier of “>1-3290398-899773.77-1-1-0” which denotes a sequence ID of 1, 3290398 raw read count, 899773.77 normalized read count, cluster ID of 1, rank within cluster of 1, and edit distance of 0 from seed. Within a given cluster ID, the seed sequence is the listed first, is the highest ranked aptamer sequence and will have an edit distance from the seed of 0. Subsequent clusters can be identified by the cluster ID. For example, in **Figure 4A**, the second cluster begins with the following identifier “>2-41052-11225.85-2-1-0”. FASTAptameR 2.0 includes the option to output a CSV data table (**Figure 4B**), which makes the clustering results sortable and easier to parse separate clusters.

**Figure 4:**
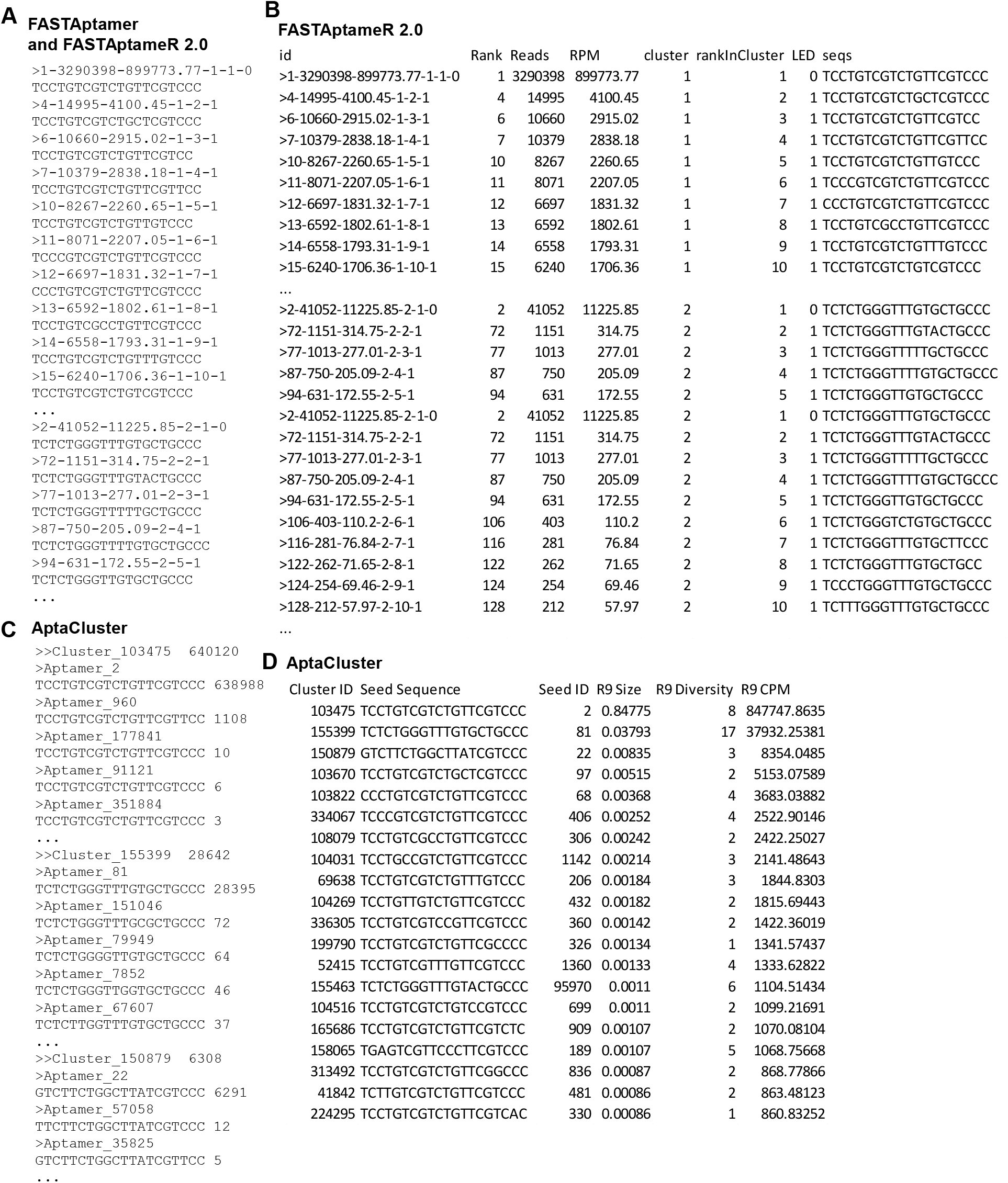
Clustering results from FASTAptamer, FASTAptameR 2.0 and AptaCluster. (A) The modified FASTA file clustering results from FASTAptamer and FASTAptameR 2.0. The FASTA header information follows as rank, reads, reads per million (RPM), cluster id, rank within the cluster, and edit distance from the seed sequence. (**B**) FASTAptameR 2.0 clustering results outputted as a data table. (**C**) AptaCluser exported clustering results from round 9 and (**D**) exported cluster table. Data shown are only a portion of larger files. Gaps in data are denoted by a “…”.

AptaCluster outputs two data files: a modified FASTA file and a data table of seed sequences (**Figure 4C, D**). The AptaCluster modified FASTA file (**Figure 4C**) includes a sequence identifier line (denoted by “>>”) with the cluster ID followed by the aptamer sequences within that cluster (denoted by “>”), starting with the seed sequence. Read counts are included on the sequence line following the aptamer variable region sequence. For example, in **Figure 4C**, the first cluster is identified as Cluster 103475, with a total of 640,120 read counts among all sequences within that cluster. The first listed sequence, which is the seed of Cluster 103475, is named Aptamer_2 and has a read count of 638,988. The AptaCluster seed sequence data table includes the cluster ID, seed sequence, and information about the clusters from each selection round; example data are provided for Round 9 in **Figure 4D**. For each selection round, AptaCluster seed table provides the proportion of a cluster relative to other identified clusters (“R9 Size”), the number of sequences within that cluster (“R9 diversity”) and the total number of normalized read counts per million (“R9 CPM”).

By comparison, AptamerRunner yields a visual output of clusters as depicted in **Figure 2A**. In this example using the same data that were analyzed by FASTAptamer, FASTAptameR 2.0 and AptaCluster, the seed sequence was the central node in the top left cluster (cluster #1). Note that all programs identified the same sequence as the seed, but the visual output by AptamerRunner significantly improves data interpretation and candidate aptamer selection because all clusters can be easily viewed simultaneously. Users can determine aptamer sequence, predicted secondary structure, and any other imported metadata (e.g., fold enrichment) by simply clicking on a node (diagrammed in **Supplemental Figure S3**). This interactive visualization is not possible with text-based results. FASTAptameR 2.0 does include functions to visualize an analysis of the clustering data within the *diversity* module. The diversity module provides a series of graphs (cluster metaplots) that depict information about the population of clustered aptamers (**Figure S4A**, sequence count, read count and average LED) and a PCA plot of a k-mer matrix (**Figure S4B and S4C**). While these PCA plots can provide insight into relative diversity of the different clusters, this function is limited to plotting no more than 15 clusters concurrently, and users must cross-reference text-based results to determine the identify of specific aptamer sequences within the plot. Results from AptamerRunner are also interactive. The limitations of FASTAptamer-Cluster, FASTAptameR 2.0 and AptaCluster text-based outputs highlight the importance of visualizing clustering data to provide a natural context for representing the different clusters of aptamers.

### Improving candidate aptamer selection by integrating fold enrichment data and scoring algorithm results onto AptamerRunner clustered aptamers

A limitation of both the text-based and visualized clustering results is that they do not provide insight into which clusters are most likely to yield the best aptamers. Clustering separates aptamers into groups that likely target the same epitope, but additional data are necessary to score and rank the aptamers to identify ideal candidates from within each cluster of aptamers. We hypothesized that integration of fold enrichment or data from scoring algorithms significantly enhances candidate selection when integrated with clustering results. Unfortunately, the text-based outputs of FASTAptamer and FASTAptameR 2.0 do not allow for easy integration of fold enrichment or scoring data. The AptaCluster seed sequence data table provides selection round normalized read counts that can be used to calculate the overall round-to-round enrichment of each cluster (see **Figure 4D**), but data from this table must be manually cross-referenced with the clustering results to determine the enrichment of all sequences within the cluster.

AptamerRunner overcomes the limitations of text-based results through visual integration of fold enrichment data and results from scoring algorithms. Data tables were imported into Cytoscape; these tables contained log10 normalized read counts of round 9 and the log2 fold enrichment between selection rounds (e.g., round 2 to round 5) of aptamers that were clustered using the seed approach with an edit distance of 1. As shown in **Figure 5**, such visual overlays of node size and color provide easy to interpret visual cues of individual aptamer abundance and enrichment during the SELEX process. Groups of aptamers showing positive enrichment (green) from negative enrichment (yellow) are easily discernable. Furthermore, the log2 fold enrichment can be easily evaluated between different selection rounds. The B-cell SELEX process sequence enrichment exhibited a sigmoidal curve, with rounds 0 to 2 representing the initial exponential phase, rounds 2 to 5 representing the linear phase and rounds 5 to 9 representing the asymptotic phase (**Figure 5A**). Interestingly, with this visual comparison we observed positive enrichment from rounds 0-2 of the selection (**Figure 5B**), but the most the most interesting changes of log2 fold enrichment seem to occur between selection rounds 2 to 5 (**Figure 5C**) and rounds 5 to 9 (**Figure 5D**). For example, when comparing round 5 to 9, the largest group of clustered aptamers (top left cluster) has a significant diversity of log2 fold enrichments that ranges from −9.95 to 5.1, and the seed sequence remained close to 0 (**Figure 5D**). These data suggest that the seed sequence may not be the ideal candidate from this cluster, but rather one of the other aptamer sequences within 1 edit distance of the seed sequence with a positive log2 fold enrichment is preferred for subsequent testing. By contrast, the seed sequence in the top right cluster is a desirable candidate based on its positive log2 fold enrichment from round 5 to round 9. This type of analysis of AptamerRunner clustering results mapped with log2 fold enrichment data was used with the Ruiz-Ciancio *et. al.* study to identify 38 candidate aptamers from 38 separate sequence clusters (all-vs-all approach with edit distance 1) or structure clusters (all-vs-all approach with tree distance 3). Within each cluster, candidates were defined as the aptamer sequences with the highest log2 fold enrichment from rounds 2 to 5 or rounds 5 to 9. We favor the all-vs-all approach based on the supposition that the seed approach may miss interesting inter-aptamer relationships; however, we present data in this head-to-head comparison by using the seed approach since other aptamer clustering tools cannot accommodate the all-vs-all approach.

**Figure 5:**
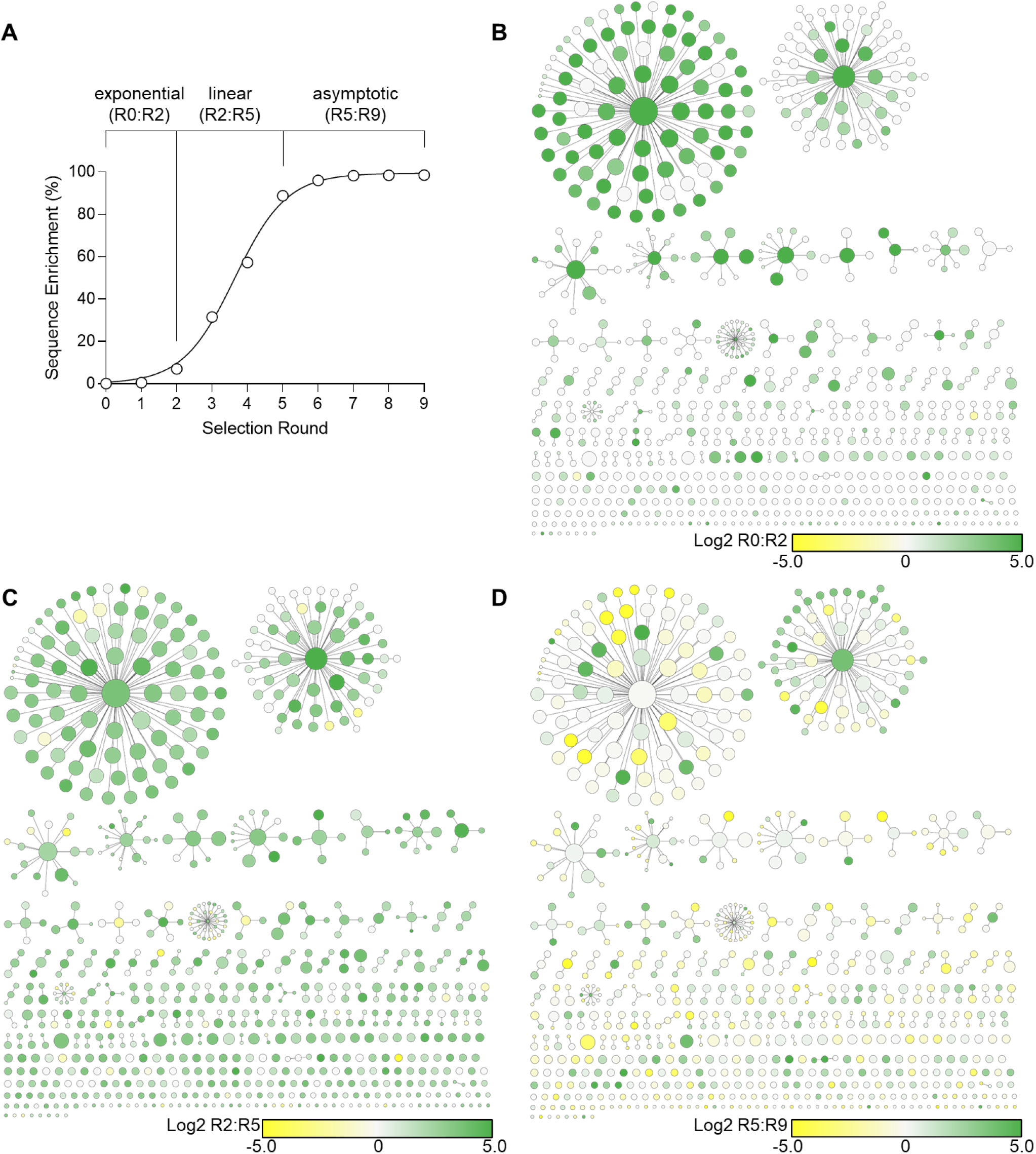
Visual integration of round-to-round log2 fold enrichment data mapped onto clustered aptamers. The log2 fold enrichment data between selection rounds from a SELEX process imported into the Cytoscape data table can be applied to determine the visual properties of aptamers clustered by AptamerRunner. The node size was set to the log10 normalized read count of round 9 and node color was determined by the log2 fold enrichment between different selection rounds based on (**A**) the phases of the sequence enrichment % sigmoidal curve fit; exponential (R0:R2), linear (R2:R5) and asymptotic (R5:R9). (**B**) The log2 fold change enrichment of round 0 to round 2 shows early linear phase enrichment of aptamer sequences, (**C**) round 2 to round 5 shows changes in aptamer sequences during the linear phase of sequence enrichment when the aptamer library experienced the greatest changes in convergence and (**D**) round 5 to round 9 show changes in aptamer sequences during the asymptotic phase when the aptamer library had reached maximum convergence.

Bioinformatics tools such as MPbind (17) and RaptRanker (16) score and rank aptamers enriched during SELEX, and these data can also be overlayed onto AptamerRunner clustering data in Cytoscape. The MPbind scoring algorithm generates a combined meta z-score of aptamer sequences based on relative motif enrichment and abundance of the final aptamer selection round. RaptRanker evaluates aptamer sequence motif and structure of subsequence groups to generate an average motif enrichment score (AME). Integrating data from scoring algorithms enables us to examine the predicted relative aptamer affinities, which can be used to select candidates within each cluster. Interestingly, MPbind and RaptRanker demonstrates significant variation in the predicted affinities of aptamers from the B-cell SELEX dataset (**Figure 6**). Specifically, MPBind predicted that most of the clusters had high affinity for their targets: the majority of the nodes were visualized as red (meta z-scores of 40-60, **Figure 6A**), whereas RaptRanker predicted fewer high affinity aptamers, with most nodes visualized in the blue to yellow spectrum (average motif enrichment of 0-5, **Figure 6B**). Since MPBind and RaptRanker use different principles for scoring, we asked if the highest scoring aptamers were similar between the two algorithms by plotting the scores as interactive scatter plots in Cytoscape. Clusters of aptamers that were scored high by both the algorithms could be identified, but, consistent with the visual representation of the clusters, many more aptamers scored highly by MPBind vs. RaptRanker (**Figure 6C**).

**Figure 6:**
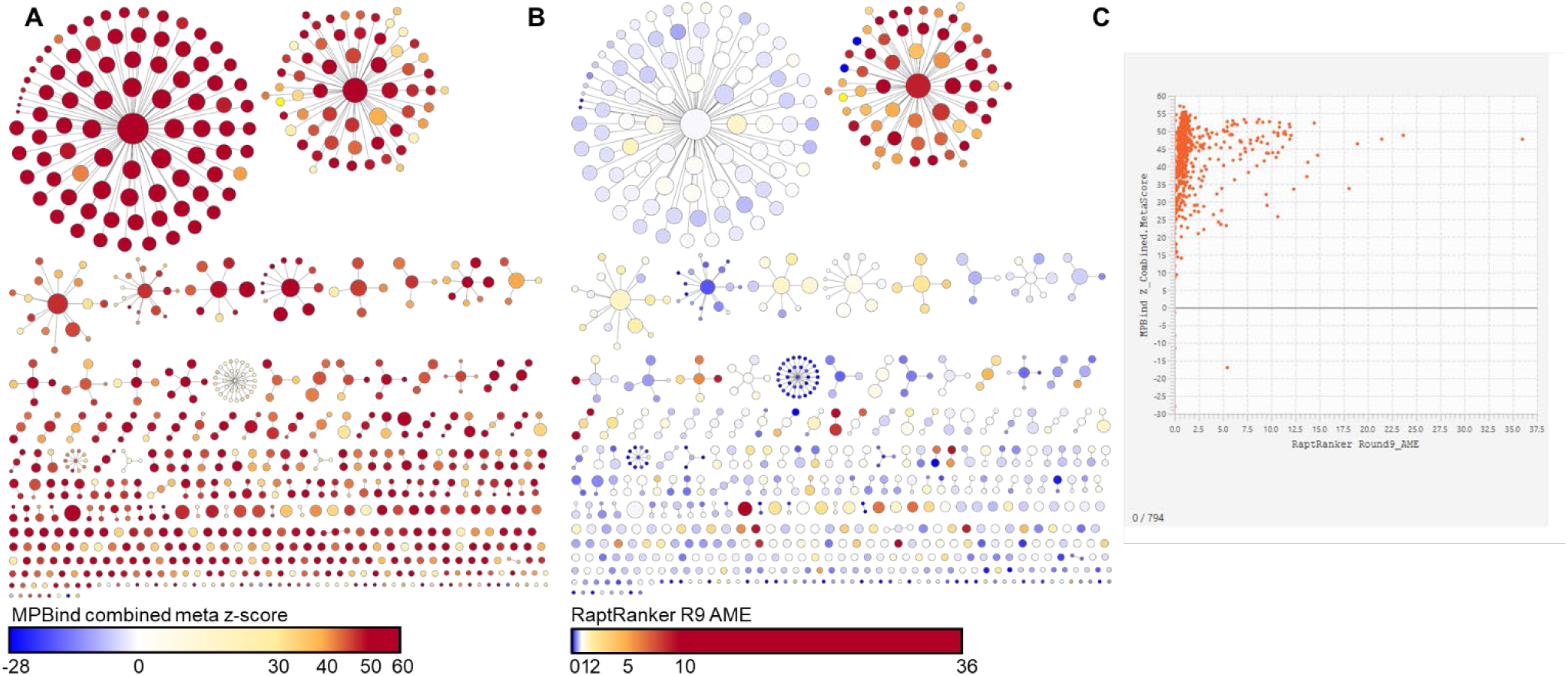
Aptamer scoring algorithm mapped to clustered aptamers. Data from aptamer scoring algorithms (**A**) MPBind and (**B**) RaptRanker were mapped to the networks generated by AptamerRunner. (**C**) Cytoscape can produce scatter plots that can compared the scoring data from MPBind and RaptRanker to identify aptamers and clusters of aptamers that were scored highly by both algorithms.

### Application of AptamerRunner metadata to re-organize networked aptamers

One rationale for using the seed approach rather than the all-vs-all approach to cluster aptamers is that the all-vs-all approach will frequently generate large, disorganized hairball clusters of aptamers as highlighted in **Figure 7A** (grey nodes and red edges), which are difficult to interpret. To address this limitation with all-vs-all data, the metadata generated by AptamerRunner during clustering can be used to deconstruct and re-organize the hairballs. For example, the hairball cluster within **Figure 7A** was isolated and the aptamer sequences re-organized using the metadata property of structure relatedness. This approach identified groups of aptamer sequences that are related by edit distance 1 and have identical structures (tree distance = 0), as shown by edges colored blue (**Figure 7B**). Several groups of the re-organized aptamer sequences also exhibited positive log2 fold enrichment from round 5 to round 9 (green nodes). We also asked if this de-convolution approach could be used for other metadata features. In the example shown in **Supplemental Figure S5**, we tested whether enriched aptamers within this hairball are structurally similar. First, we re-organized the hairball in **Supplemental Figure S5A** using the log2 fold change enrichment observed between round 5 to round 9 (Log2 R5:R9, **Supplemental Figure S5B**). Next, we excluded any aptamers with a log2 R5:R9 less than ≤ 0, and the remaining subset of aptamers were reorganized for identical predicted structures (tree distance = 0, **Supplemental S5C**). This approach identified multiple sub-groups of aptamer sequences within the hairball network that were positively enriched during SELEX and are closely related by both sequence and structure. Taken together, these capabilities of AptamerRunner to incorporate metadata for clusters to be visualized in Cytoscape allow for a more in-depth analysis and identification of candidate aptamers. Use of different parameters provides greater resolution of the enriched sub-groups of aptamers, which is a major improvement by AptamerRunner over text-based clustering algorithms.

**Figure 7:**
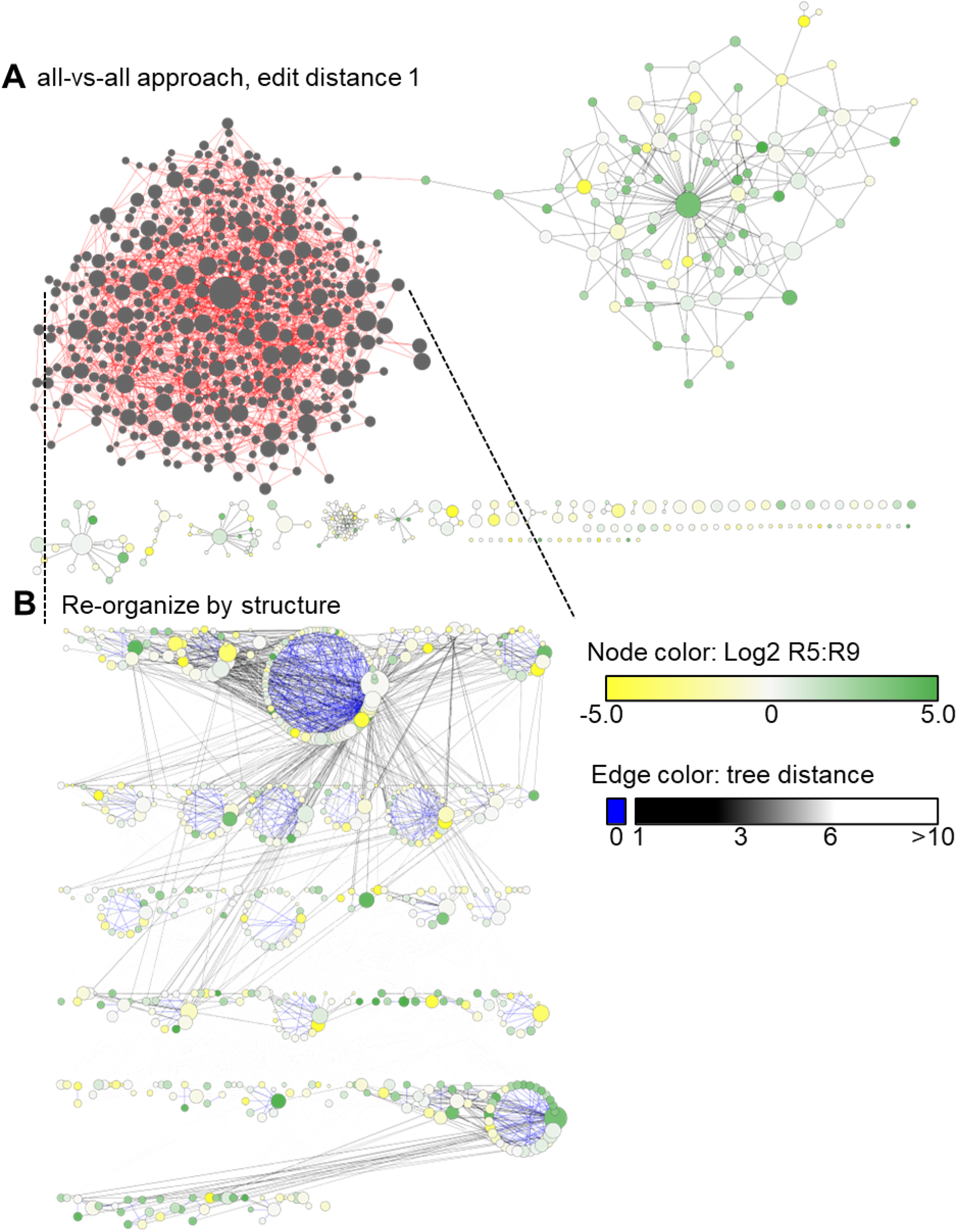
Re-clustering aptamer networks within Cytoscape. Groups of aptamers can be re-clustered using tools within Cytoscape. (**A**) Nodes within a large hairball network of aptamers clustered by edit distance 1 using the all-vs-all approach were selected and (**B**) re-organized within Cytoscape by identical structures.

## DISCUSSION

The idea behind clustering aptamer sequences is that groups of aptamers related by sequence and structure likely target the same epitope and conversely, groups of aptamers not related likely target different epitopes. Therefore, understanding how aptamers are related, and not related, is critical for identifying a diverse cohort of aptamers for experimental validation. With the AptamerRunner clustering algorithm, distance measures are now retained as metadata and the option to use logical operators (AND, OR) for clustering was introduced. Incorporating distance measures as metadata within the edges provides additional opportunities to analyze, re-evaluate and interpret clustering results. Within Cytoscape, additional data such as the log2 fold enrichment of aptamer sequence across selection rounds and data from other aptamer bioinformatics scoring algorithms can be mapped onto the networks generated by AptamerRunner to aid in the interpretation of the clustering results. Having the AptamerRunner Docker container controlled by a .NET program presents a simple method to enable users to access AptamerRunner. Users only need to use a command line interface to initiate the .NET script, which then dynamically generates all Docker commands necessary to run AptamerRunner’s secondary structure prediction or clustering algorithms. Taken together, the innovative features of AptamerRunner enable a more in-depth analysis of aptamer NGS data and allows for better identification of candidate aptamers. In addition, AptamerRunner can be used to ask new questions about how sequence and structure relatedness contribute to library convergence during the SELEX process.

The superiority and flexibility of AptamerRunner is highlighted by two recent publications that made use of AptamerRunner clustering capabilities in fundamentally different ways (13,14). Ruiz-Ciancio *et. al.* (14), using the same NGS dataset as in the present study, applied the AptamerRunner clustering algorithm to identify 38 candidate aptamers from unique clusters related by sequence or by structure. These 38 candidates were then ranked for their potential to bind the CD22 protein through a molecular docking and molecular dynamic approach (14). The aptamer that exhibited specificity for CD22, B-ALL1, was the 573^rd^ ranked aptamer and was identified within a group of aptamers related by structure. The B-ALL1 aptamer would not have been identified using only an edit distance clustering approach. This highlights the importance of clustering by related structures, which is not an available capability in FASTAptamer, FASTApameR 2.0 or AptaCluster.

A second study by Santana-Viera *et. al.* (13) applied AptamerRunner to examine the relatedness of aptamer libraries enriched in two independent SELEX processes: a protein-based SELEX using recombinant human EphA2 as target and a Cell-Internalization SELEX using EphA2-expressing MDA231 cells as targets (13,37). The AptamerRunner clustering algorithm identified a group of aptamers within these two different SELEX processes that shared structure and sequence relatedness. From this group of aptamers, the candidate aptamer ATOP, was observed to target hEphA2 and exhibited antitumorigenic effects *in vitro* and *in vivo*. The flexibility of AptamerRunner permitted the researchers to develop a novel clustering strategy that made use of both edit distance and tree distance clustering using the all-vs-all approach. A seed approach with two different SELEX processes would have been challenging due to the complication in defining which SELEX would provide the aptamer sequences to serve as seeds. Importantly, without the in-depth analysis of aptamer relatedness enabled by AptamerRunner, the aptamers described by these two studies would have been prohibitively challenging to identify with the other available clustering tools.

The aforementioned examples of AptamerRunner identifying candidate aptamers made use of the all-vs-all clustering approach rather than the seed approach. The seed approach, introduced by AptaCluster and FASTAptamer, is founded on the idea that certain aptamer sequences, called the seed, serve as the basis from which mutations during SELEX accumulate to yield more specific or higher affinity aptamers. However, the seed approach for clustering aptamers is a greedy process whereby sequences connected to the seed are removed from the pool of aptamers available for clustering. Therefore, the seed approach will miss inter-aptamer connections identified by the all-vs-all approach. However, the all-vs-all approach is a significantly more computationally intensive process than the seed approach and can yield hairball networks that are more complex to interpret. The concept of the seed sequence is potentially more important for aptamer libraries with longer variable regions. Longer variable region libraries (e.g. >30 nucleotides) have a large starting complexity that cannot be sampled at the start of SELEX and PCR-generated mutations are more likely to introduce beneficial aptamer sequences not present during the initial selection rounds. FASTAptameR 2.0 includes a function (distance module) that can specifically evaluate edit distance from a seed sequence, or other sequences, to evaluate accumulation of mutations during SELEX. The FASTAptameR 2.0 distance module plots the edit distance distribution from the seed sequence as a histogram. Given that the accumulation of mutations is more likely to occur with more abundant aptamers and less likely with aptamers of lower abundance, AptamerRunner could interrogate larger more complex networks of related aptamers identified by the all-vs-all approach by re-evaluating them using the seed approach. While AptamerRunner does not intend to settle this debate, it provides the necessary tools that will allow aptamer researchers to investigate their data independently based on their goals and how the clustering data should be visualized.

Future versions of AptamerRunner could include additional aptamer bioinformatics tools to process, compile, and analyze raw aptamer NGS data with a GUI interface like the integrated pipelines offered by FASTAptameR 2.0 and AptaCluster. The structure prediction algorithm could incorporate additional structure prediction algorithms such as Mfold (38) or be modified to permit multiple structures for each aptamer sequence including suboptimal structures. Furthermore, AptamerRunner does not include any option to limit what aptamers are clustered. AptaCluster applies a hashing function that filters the dataset by identifying pairs of aptamers that are likely to be dissimilar and FASTAptameR 2.0 includes a filter function to cluster only aptamer sequences of a minimum abundance or produce a set number of clusters. With AptamerRunner we filtered the aptamer NGS dataset prior to clustering using a separate aptamer abundance and persistence analysis (14,39). Ideally, algorithms that can filter aptamer NGS datasets, such as the AptaCluster hashing function, could be applied independently prior to clustering to investigate the effectiveness of different filtering strategies. Additional future directions include determining the range between separate groups of networked aptamers with the idea that more distantly related aptamers are more likely to target different epitopes. Analysis options could include ranking different groups of clusters aptamers and ranking individual aptamers within each cluster by integrating scoring or application of molecular docking to predict which groups of aptamer bind to same regions of a target protein (17,36,40-43).

## CONCLUSION

In summary, AptamerRunner stands out as a powerful and versatile tool for clustering aptamer NGS data with innovative approaches, enabling diverse sequence and structure relatedness, introducing logical operators, and offering seamless integration with Cytoscape for visualization and interpretation. The inherent flexibility, user-friendly design and prospects for future enhancements collectively position AptamerRunner at the forefront of advancing aptamer research.

## DATA AVAILABILITY

Source data (14) are provided with this paper in supplemental. Further data supporting the findings of this study are available from the corresponding author upon reasonable request. The AptamerRunner .NET script Windows and Linux versions are available in supplemental and on GitHub (https://github.com/ui-icts/aptamer-runner/releases/tag/v0.0.3).

## SUPPLEMENTARY DATA

Supplementary data are available online at NAR.

## AUTHOR CONTRIBUTIONS

Dario Ruiz-Ciancio: Investigation, Formal Analysis, Visualization, Writing – original draft. Suresh Veeramani: Investigation, Formal Analysis, Writing – review & editing. Eric Embree: Software, Resources, Data curation. Chris Ortman: Software, Resources, Supervision. Kristina W. Thiel: Formal Analysis, Writing – original draft. William H. Thiel: Conceptualization, Methodology, Formal Analysis, Project administration, Writing – original draft.

## SUPPLEMENTAL FIGURES

**Figure S1:**
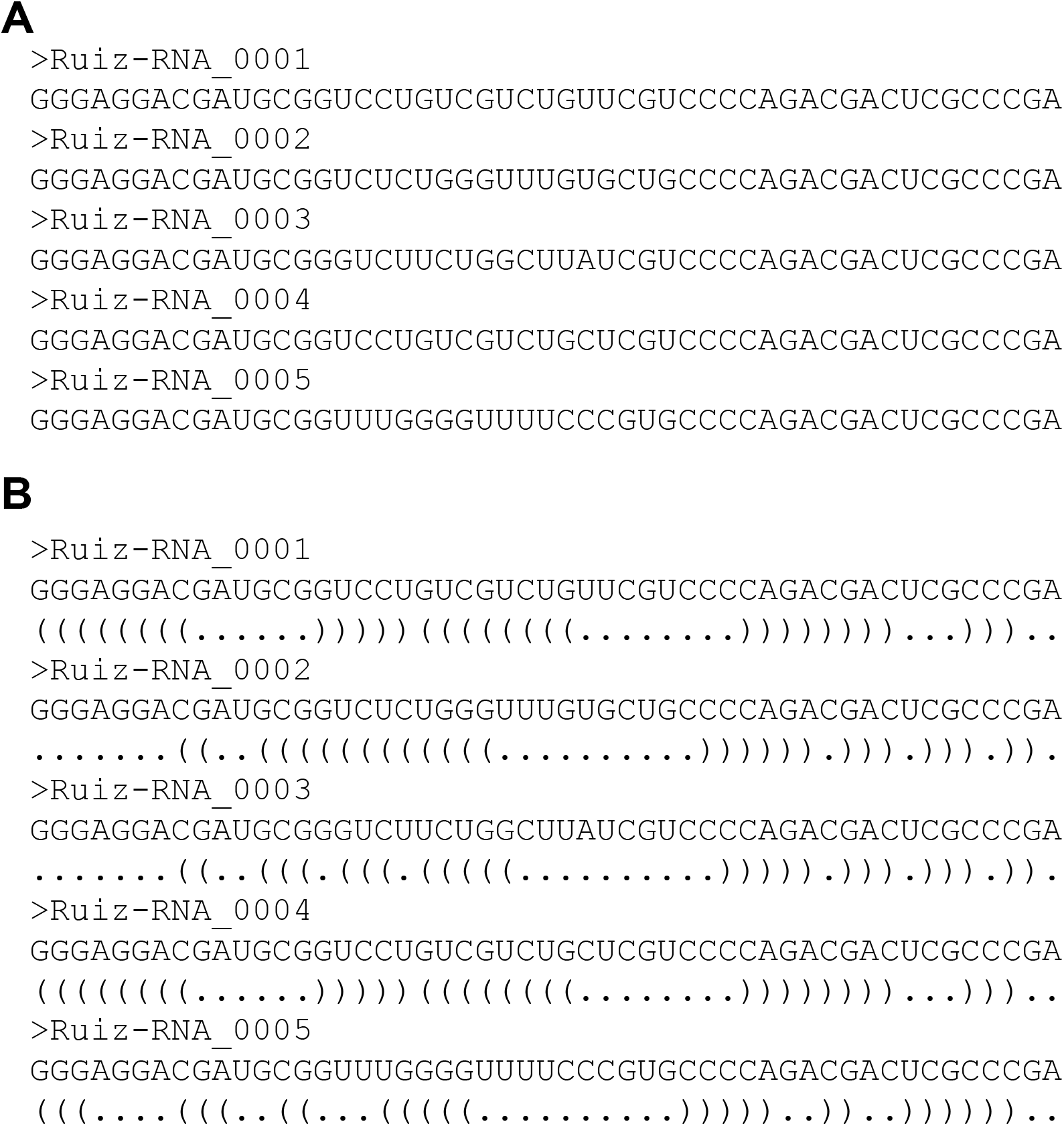
Aptamer sequences within a FASTA-formatted file. (**A**) The AptamerRunner structure prediction algorithm requires a FASTA-formatted file (e.g. >Ruiz-RNA_0001). Within the FASTA file, the aptamer sequences are ranked in descending order with the most abundant aptamer listed first. (**B**) The modified FASTA-formatted file outputted by the AptamerRunner structure prediction algorithm (FASTA.struct) includes the predicted aptamer secondary structures in dot-bracket annotation. The secondary structure of each full-length aptamer sequence is predicted using the RNAfold (1) structure prediction algorithm from the Vienna Package (v2.0) (2).

**Figure S2:**
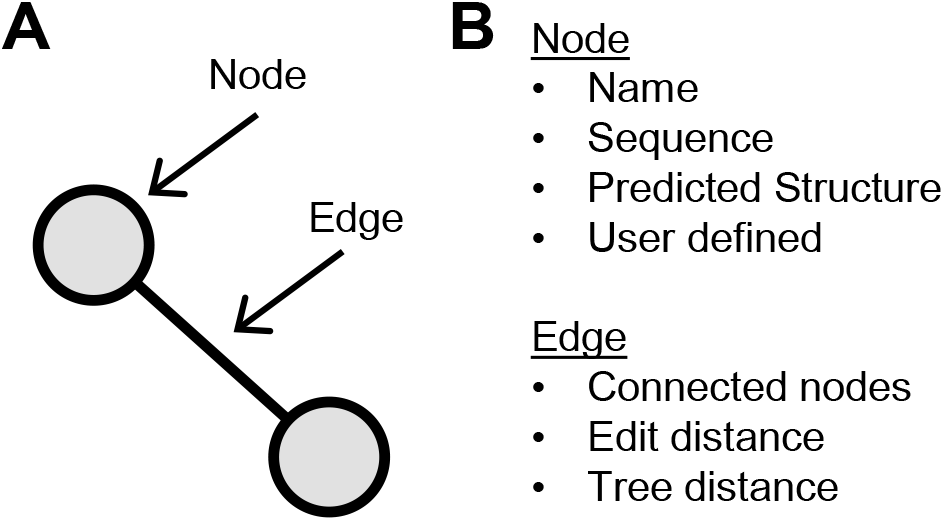
Cytoscape visualization of interconnected aptamers. AptamerRunner clustering results are visualized within Cytoscape using nodes, representing unique aptamer sequences; and edges that connect related aptamer sequence nodes by a set distance measure (e.g. edit distance 1). Nodes are associated with metadata that include the aptamer name, sequence, and predicted structure); edges comprise metadata associated with the edit and tree distance measures between two connected nodes.

**Figure S3:**
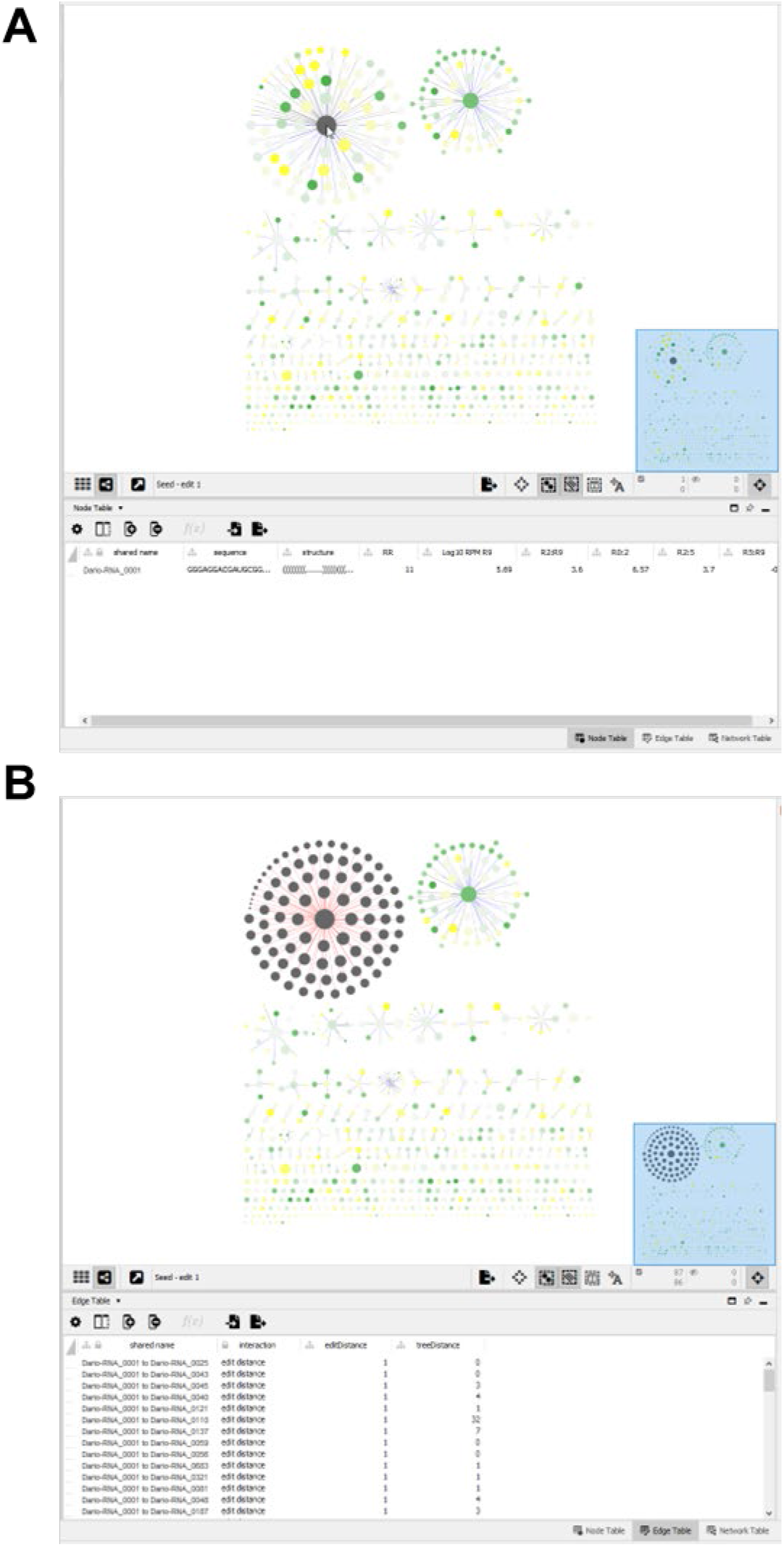
Cystoscope’s interface flexibility. (A) Cystoscope’s interactive interface permits easy and precise selection of nodes. Selected nodes (cursor with single node selected in grey) with all corresponding node data of each aptamer can be viewed in the node table below, which may be copied directly or exported as text files. (**B**) AptamerRunner distance measure data of interconnected aptamers (cluster with grey nodes and red edges) is found within the Edge Table of Cytoscape.

**Figure S4:**
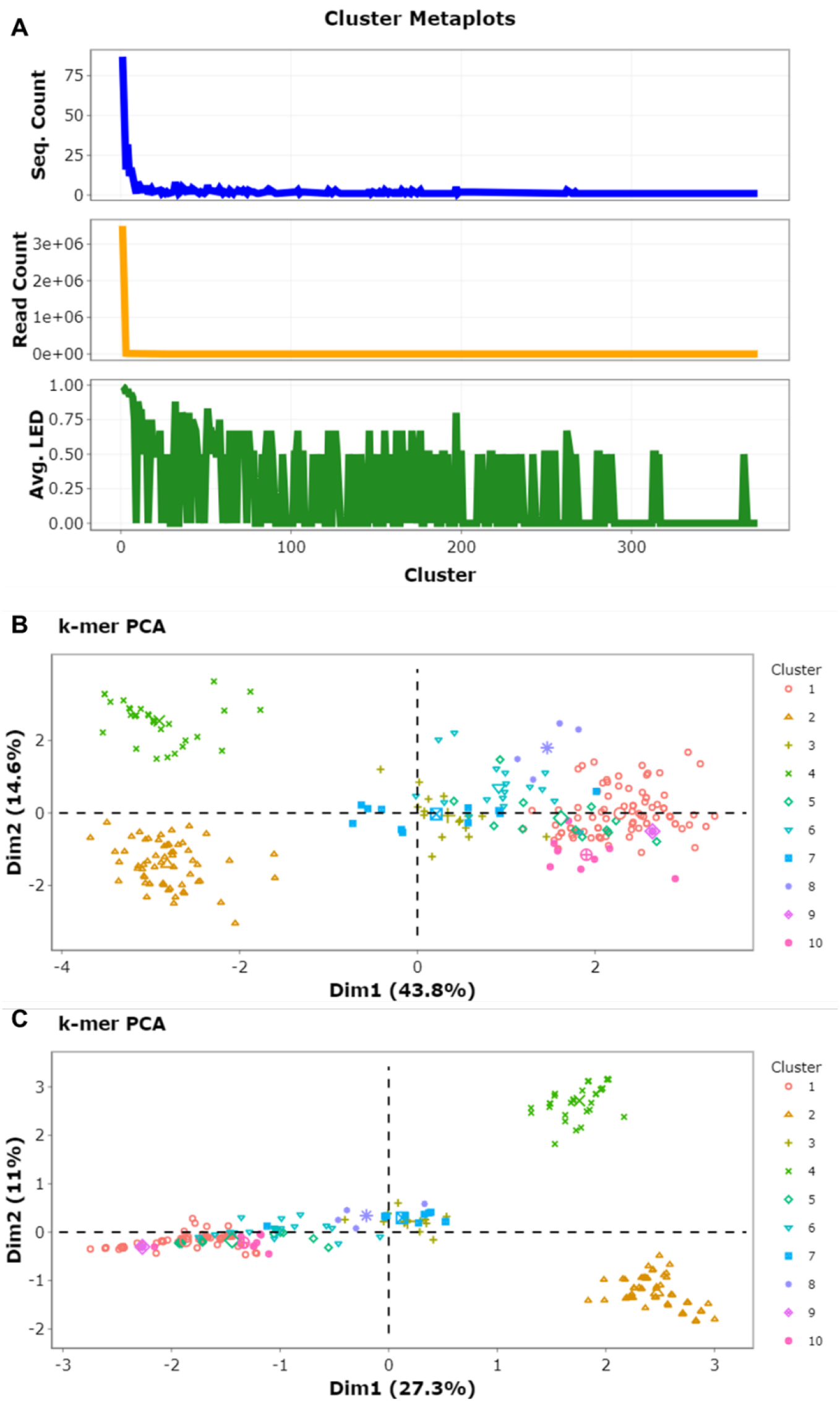
FASTAptameR 2.0 diversity module plots. (**A**) Meta plots. (**B**) K-mer 3 PCA plot of clusters 1-10. (**C**) K-mer 5 PCA plot of clusters 1-10.

**Figure S5:**
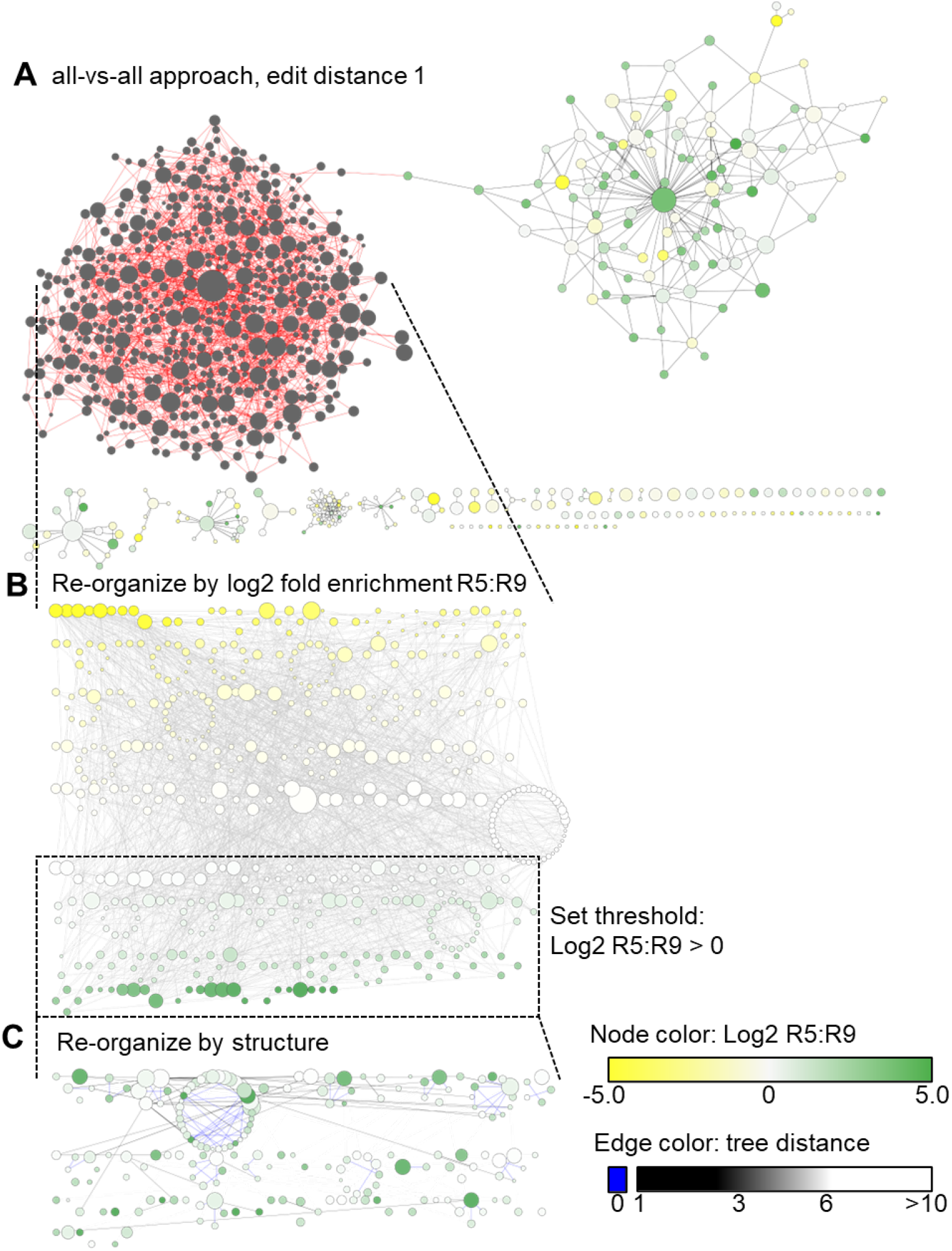
Re-organization of networks using Cytoscape. (**A**) The large cluster of aptamers generated in Figure 7 (figure above, highlighted as grey nodes and red edges) was (**B**) re-organized by the log2 fold enrichment between round 5 and round 9 (Log2 R5:R9). Groups of aptamers with log2 fold enrichment R2:R9 greater than 0 were selected and (**C**) re-organized by structure relatedness.

## SUPPLEMENTAL METHODS

### NGS aptamer dataset

The aptamer NGS dataset from Ruiz *et. al* (3). was used to demonstrate AptamerRunner capabilities. The dataset consisted of enriched libraries of a Cell-Internalization SELEX against human B-cell acute lymphoblastic leukemia cell lines. The FASTQ data were processed into a non-redundant database (NrD) using Galaxy workflows (4). The NrD was analyzed for aptamer abundance and aptamer persistence and then filtered to 794 sequences (supplemental data) for clustering by AptamerRunner. The same dataset was used for running other clustering and affinity scoring tools mentioned in this manuscript.

### Cytoscape

Cytoscape version 3.7.2 was used to display clustering XGMML results. Data were imported into Cytoscape using *File\Import\Network from file…* and displayed using *Layout\yFiles Organic Layout*. Properties of aptamer sequences were compiled using Microsoft Excel and included aptamer name, log10 normalized selection round read counts per million, log2 fold enrichment between different selection rounds and dot-bracket predicted aptamer structure. These data were exported as either comma-delimited or tab-delimited text files and imported into Cytoscape using *File\Import\Table from File…* using the aptamer name as the key. Node shape, size and color were modified using Cytoscape *Style* options based upon the imported network properties. Interconnected clusters of related aptamers were isolated using the Cytoscape function *Tools\Subnetwork Creation\Extract Connected Components.* Isolated aptamer clusters were re-clustered using the Cytoscape function *Layout\Group Attributes Layout* and selecting the imported aptamer dot-bracket structure property. Isolated aptamer clusters were identified using the Cytoscape function *Tools\Plott Scatter* and selecting the imported aptamer data within the *CyChart* to create the desired Scatter Plot.

### FASTAptamer-Cluster

A FASTAptamer counted formatted-FASTA file was generated from the NGS aptamer data set and used as the input file for the FASTAptamer-Cluster Perl program (5). The FASTAptamer-Cluster program was run with edit distance 1.

### FASTAptameR 2.0

The cluster module and diversity module of FASTAptameR 2.0 (6) were run using the same counted FASTA input file as used with FASTAptamer-Cluster. The FASTAptameR 2.0 cluster module was run with edit distance 1 with the number of clusters set to maximum.

### AptaCluster

AptaCluser was accessed through AptaGUI (7) using data generated by AptaPlex (8) processing FASTQ data of selection rounds 5, 7 and 9. AptaCluster was run with the default values for the k-mer hashing function and using an edit distance of 1.

### MPBind

FASTQ data of selection round 0 through 9 and a control round were processed (MPBind_Preprocess.py) and analyzed (MPBind_Train.py) using the MPBind pipeline (9). The aptamer sequences within the NGS aptamer data set were analyzed by MPBind (MPBind_Predict.py) and the combined meta z-scores of these aptamers were imported into the Cytoscape node data table.

### RaptRanker

FASTQ data of selection rounds 5, 7 and 9 were run through RaptRanker (10). The scores of the NGS aptamer data set were pulled from the RaptRanker score table and imported into Cytoscape node data table.

### Pre-requisites to run AptamerRunner

- Prerequisite software to install

- Docker or Docker Desktop
- Cytoscape
- AptamerRunner v0.0.3 .NET script (Windows or Linux version)

- Supplemental information
- https://github.com/ui-icts/aptamer-runner/releases/tag/v0.0.3

### Using AptamerRunner for the first time

- Start Docker or Docker Desktop (it could be installed from www.hub.docker.com)
- Open a command line terminal
- Run the AptamerRunner .NET script with the help function AptamerRunner help
- AptamerRunner will download the most recent AptamerRunner Docker image and display the help content.

### Using AptamerRunner structure prediction program

- Start Docker or Docker Desktop
- Open command line terminal
- Run the AptamerRunner .NET script with the “predict-structures” command, pointing to where the data file is located, any structure prediction options and where the output should be deposited.
- AptamerRunner will run the Docker image with any specified commands, output any data files as specified and then shut down the Docker image.

#### Examples

- Specifying the location of the input FASTA file

- Using relative location of the input FASTA file AptamerRunner predict-structures ./Ruiz/Ruiz-RNA_794
- Using absolute location of the input FASTA file AptamerRunner predict-structures Users/Ruiz/Ruiz-RNA_794
- Adding a 5’ and 3’ constant region AptamerRunner predict-structures ./Ruiz/Ruiz-RNA_794_no_constants --prefix “GGGAGGACGAUGCGG” --suffix “CAGACGACUCGCCCGA”
- Specifying pass through options for RNAfold(1,2) AptamerRunner predict-structures ./Ruiz/Ruiz-RNA_794 --pass_options "-p -d2 -T 25"
- Specifying the location of the output files AptamerRunner predict-structures ./Ruiz/Ruiz-RNA_794 -o ./Ruiz/Predict-Structures

### Using the AptamerRunner clustering program

- Start Docker or Docker Desktop
- Open command line terminal
- Run the AptamerRunner .NET script with the “create-graph” command, pointing to where the predicted structure data file (.struct.fa) is located, clustering options and where the output should be deposited.
- AptamerRunner will run the Docker image with any specified commands, output any data files as specified and then shut down the Docker image.

#### Examples

- *Seed* approach using edit distance 1 (Figure 2A) AptamerRunner create-graph ./Ruiz/Ruiz-RNA_794.struct.fa -t tree -d 0 --seed
- *Seed* approach using tree distance 0 (Figure 2B) AptamerRunner create-graph ./Ruiz/Ruiz-RNA_794.struct.fa -t tree -d 0 --seed
- *All-vs-all* approach using edit distance 1 (Figure 2C) AptamerRunner create-graph ./Ruiz/Ruiz-RNA_794.struct.fa -t edit -e 1
- *All-vs-all* approach using tree distance 0 (Figure 2D) AptamerRunner create-graph ./Ruiz/Ruiz-RNA_794.struct.fa -t tree -d 0
- *Seed* approach using edit distance 1 AND tree distance 0 (Figure 3A) AptamerRunner create-graph ./Ruiz/Ruiz-RNA_794.struct.fa -t and -e 1 -d 0 --seed
- *All-vs-all* approach using edit distance 1 AND tree distance 0 (Figure 3B) AptamerRunner create-graph ./Ruiz/Ruiz-RNA_794.struct.fa -t and -e 1 -d 0
- *Seed* approach using edit distance 1 OR tree distance 0 (Figure 3C) AptamerRunner create-graph ./Ruiz/Ruiz-RNA_794.struct.fa -t both -e 1 -d 0 --seed
- *All-vs-all* approach using edit distance 1 OR tree distance 0 (Figure 3D) AptamerRunner create-graph ./Ruiz/Ruiz-RNA_794.struct.fa -t both -e 1 -d 0

### AptamerRunner error

C:\User\AptamerRunner>AptamerRunner

Docker version 20.10.20, build 9fdeb9c

Using docker image ghcr.io/ui-icts/thiel-aptamer:latest

Pulling docker image to verify the latest version is present.

error during connect: This error may indicate that the docker daemon is not running.: Post “http://%2F%2F.%2Fpipe%2Fdocker_engine/v1.24/images/create?fromImage=ghcr.io%2Fui-icts%2Fthiel-aptamer&tag=latest”: open //./pipe/docker_engine: The system cannot find the file specified.

Failed to pull docker image ghcr.io/ui-icts/thiel-aptamer:latest

- If this error is encountered, Docker is not running. Start Docker or Docker desktop and re-run the AptamerRunner .NET script.

### Importing clustering results (.xggml file) into Cytoscape

- Within Cytoscape select File/Import/Network from File…
- Select the .xgmml file of the AptamerRunner clustering results
- Nodes and edges will be displayed on top of each other. Select a layout under Layout (e.g. yFiles Organic Layout was used for Figure 2 and Figure 3 clustering results) to display networks of connected aptamers.

### Importing data tables (e.g. .properties file from predict-structures) into Cytoscape

- With clustering results displayed and the node table tab selected
- In Cytoscape select File/Import/Table from File…
- Select the data file to be imported
- Use the node name as the key

